# Identification of Patient Trajectories in Timeseries Clinical Transcriptomics Data

**DOI:** 10.64898/2025.12.04.692308

**Authors:** Yixin Chen, Peyman Passban, Mahesh Raundhal, Lu Zhang, David Habiel, Sachin Mathur

## Abstract

In clinical trials, it is common for only a subset of patients to respond to a given therapy. Such variability may arise from genetic differences, environmental influences, or the presence of distinct disease endotypes. Longitudinal transcriptomic profiles collected in phase 2a/2b studies provide a unique opportunity not only to investigate drug-induced biological mechanisms in humans, but also to understand why certain individuals fail to respond and to uncover previously unrecognized disease endotypes—ultimately informing the development of targeted therapeutics. However, analyzing these datasets is challenged by substantial patient heterogeneity, variability in disease severity at each visit, and the coarse temporal resolution due to sparse sampling.

To address these limitations, we introduce a classification-guided autoencoder framework that jointly optimizes gene-expression reconstruction and classification objective to learn disease-relevant sample embeddings. Sample embeddings from all patients are leveraged to establish a continuous representation of disease dynamics, which can be clustered to delineate discrete disease states. We then construct a patient-sample graph in the learned latent space and apply a multi-commodity-flow based algorithm to infer patient trajectories through these states, enabling the identification of different patient trajectories. We evaluated our approach on three interventional datasets—ulcerative colitis, psoriasis, and atopic dermatitis, each containing both responders and non-responders. The method recapitulates known pathway perturbations associated with anti-IL17 and anti-IL6 therapies, identifies intermediate states reflecting disease/treatment progression, and reveals biologically meaningful patient trajectories within the patient population.

Code and data are available in a public GitHub repository - https://github.com/Sanofi-Public/EndotypeDetection-Timeseries

## 1 Introduction

Clinical timeseries datasets with interventions (drug treatment) offer an opportunity to uncover biological mechanisms associated with drug treatment and disease progression. These are usually phase-2a/b studies in which a small number of patients are dosed with the drug and longitudinally followed up with measurements for primary/secondary endpoints to determine efficacy of the drug. Typically, not all patients respond uniformly to drug treatment with some meeting the criteria of disease remission, while others fall short [10]. It is important to understand the reason for non-response of drug treatment in the subset of patients (non-responders) so that better patient selection can be performed in larger phase-3 trials and importantly it offers the opportunity to develop new drugs for the non-responding patients. Non-response can be attributed to different demographics, genetics, environmental factors, varied disease states and/or presence of different disease endotypes (subtypes) [15].

While the longitudinal nature of data offers several advantages over single timepoint datasets, they come with a unique set of challenges. Firstly, in clinical trials only a few timepoints are available that limit reconstruction of disease/treatment mechanisms over time. Typically patients are in different stages of the disease and if patient samples are ordered correctly [14] then samples of different patients can be leveraged to uncover dynamics of disease progression. Secondly, clinical bulk-transcriptomic datasets have increased noise levels in gene expression. While expression measures from model organisms have less variance at a given timepoint, human samples typically have large heterogeneity owing to demographics, genetics, but also due to non-uniform disease states of patients at a given visit. It is important to take into consideration the disease state of a patient at a given timepoint and not to group them with samples of other patients at the same visit who may be in different disease states. Thirdly, only a subset of genes are induced as a result of drug treatment and it is important to extract important features from a patient’s sample while constructing disease states though which a patient traverses during the course of the treatment. Clustering samples of patients in similar state is often accomplished by computing the average distance between gene expression profiles using a distance metric [1], but it may not be suitable in scenarios where only a small subset of genes change as the large gene set can dilute contribution from a small set of genes.

MANAclust [20] is an integrative framework that combined clinical data with multi-omics to identify disease endotypes using merged affinity network association clustering. The approach is for static snapshots and does not explicitly model the temporal disease progression. Truffle [8] models time-series data by first clustering samples and then applies a multi-commodity flow (MCF) problem solver to model transitions. While Truffle uses a powerful network flow concept that utilizes intermediate disease states, it uses PCA to model linear relationships and uses average euclidean distance between clusters. These approaches are not suitable to capture non-linear relationships and tend to dilute gene expression in cases where a small set of genes drive the drug response, which is often the case. Approaches used to model trajectories in timeseries single cell data such as Tempora [19] and psupertime [12] can be adapted for bulk datasets to extract potential patient trajectories. However, they have hard constraints on use of sample collection times which can result in high variance due to varied disease states among patients. Furthermore, these approaches are designed for single cells that typically have thousands of cells compared to *<* 50 samples in bulk datasets.

Variational Autoencoder (VAE) [9] has been used for non-linear dimensionality reduction for gene expression studies. As demonstrated in [13], VAEs significantly outperform traditional linear methods such as PCA and NMF in settings of high-dimension and low-sample size (*p ≫ N* ) by capturing complex data structures, which leads to superior performance in downstream unsupervised clustering tasks. Researchers in [7] demonstrated that creating low-dimensional embeddings of complex molecular features with a VAE results in significantly more stable and meaningful clusters compared to applying clustering algorithms directly on the original high-dimensional data.

In this study, we address the issue of representation of samples of patients to construct meaningful disease states by using a VAE for dimensionality reduction. We combine the reconstruction loss with the classification loss to make representations distinguishable between healthy/pretreatment and treatment. The goal of auxiliary loss is to inject domain knowledge in to the network so that the sample embeddings learn the local context of the dataset. The representations not only capture non-linear relationships among genes, but also help in overcoming the noise in the datasets. The sample embeddings are then used to construct meaningful disease/treatment states. We then make use of multi-commodity flow [17] problem solver to compute the trajectory of each patient across disease states to finally obtain major patient trajectories associated with the drug treatment. We tested the method on 3 intervention studies that were associated with phase-2 trials in ulcerative colitis (UC), atopic dermatitis (AD) and psoriasis. Results uncover mechanisms that are regulated by the intervention such as pathways regulated by anti-IL17, anti-IL6 treatments, confirm previously known findings in literature, uncover intermediate disease states that reflect disease/treatment progression and show interesting trajectories within the patient population. Lastly, we compared the method with Truffle and show the comparative results on the 3 datasets.

## 2 Data and Methods

In this section, we describe the data and methods we used in the study.

### 2.1 Datasets

We applied the method on 3 intervention datasets from phase-2 clinical trials. All datasets were downloaded from NCBI-GEO. Table 1 shows the datasets, tissue type, intervention, and number of patients.

**Table 1.**
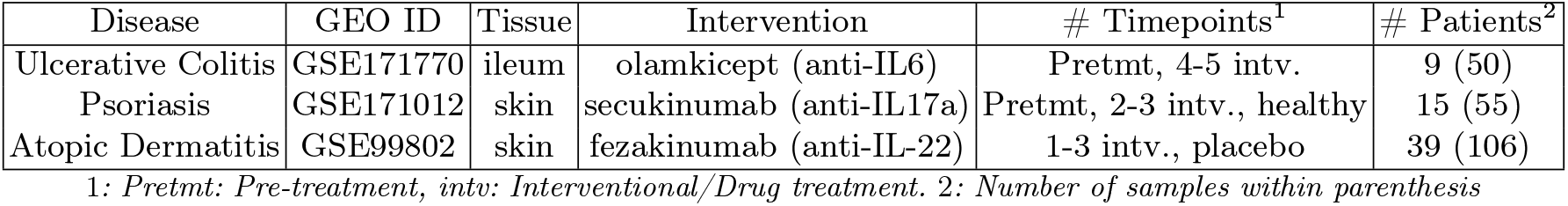
Dataset Description.

| Disease | GEO ID | Tissue | Intervention | # Timepoints<sup>1</sup> | # Patients<sup>2</sup> |
| --- | --- | --- | --- | --- | --- |
| Ulcerative Colitis | GSE171770 | ileum | olamkicept (anti-IL6) | Pretmt, 4-5 intv. | 9 (50) |
| Psoriasis | GSE171012 | skin | secukinumab (anti-IL17a) | Pretmt, 2-3 intv., healthy | 15 (55) |
| Atopic Dermatitis | GSE99802 | skin | fezakinumab (anti-IL-22) | 1-3 intv., placebo | 39 (106) |

1: Pretmt: Pre-treatment, intv: Interventional/Drug treatment. 2: Number of samples within parenthesis

In the psoriasis dataset [11], 15 patients were treated with secukinumab, an IL-17a inhibitor and samples were collected at pre-treatment, 2 weeks, 4 weeks, and 12 weeks. The PASI (Psoriasis Area Severity Index) clinical scores were used to determine that all patients had a progressive reduction in severity of the disease and all patients were deemed as responders. Patients in the UC [16] and AD [3] studies variably responded to the intervention, with some designated as non-responders in the UC study based on the clinical score (Mayo). The AD study reported that the response to the intervention depended on the expression level of IL-22 (microarray and RT-PCR), where patients with high levels of expression being responders while others were non-responders. The patient identifiers for response and non-response were not given in the dataset. In psoriasis healthy samples were used as controls, while pretreatment samples were used as controls in UC for performing GSEA. In case of AD, samples from patients that were administered placebo were used as controls. Some of the patients had data available for only a few timepoints and had missing data for the later timepoints. We used the samples of such patients in the analysis, but did not construct disease trajectories for the lack of data.

### 2.2 Data Preprocessing and Filtering

Raw gene counts were obtained from the supplementary files in NCBI GEO for the two RNASeq datasets (Psoriasis and UC). Only protein-coding genes that had at least 5 counts in at least 1% of the samples were kept. In the case of duplicate gene identifiers, the gene with highest mean expression was retained. The datasets were then normalized using TMM [4]. This was used as an input to ComBat [23] batch correction to obtain batch corrected expression values. The AD dataset was based on microarray gene expression. The normalized values for gene expression were available part of the dataset in NCBI-GEO.

After pre-processing, the expression matrix was filtered to retain the top 2,000 genes with the highest Interquartile Range (IQR), selecting features with significant variability while maintaining robustness against outliers (Figure 1a). Several other cutoffs of up to 5000 genes were also tested.

**Fig. 1.**
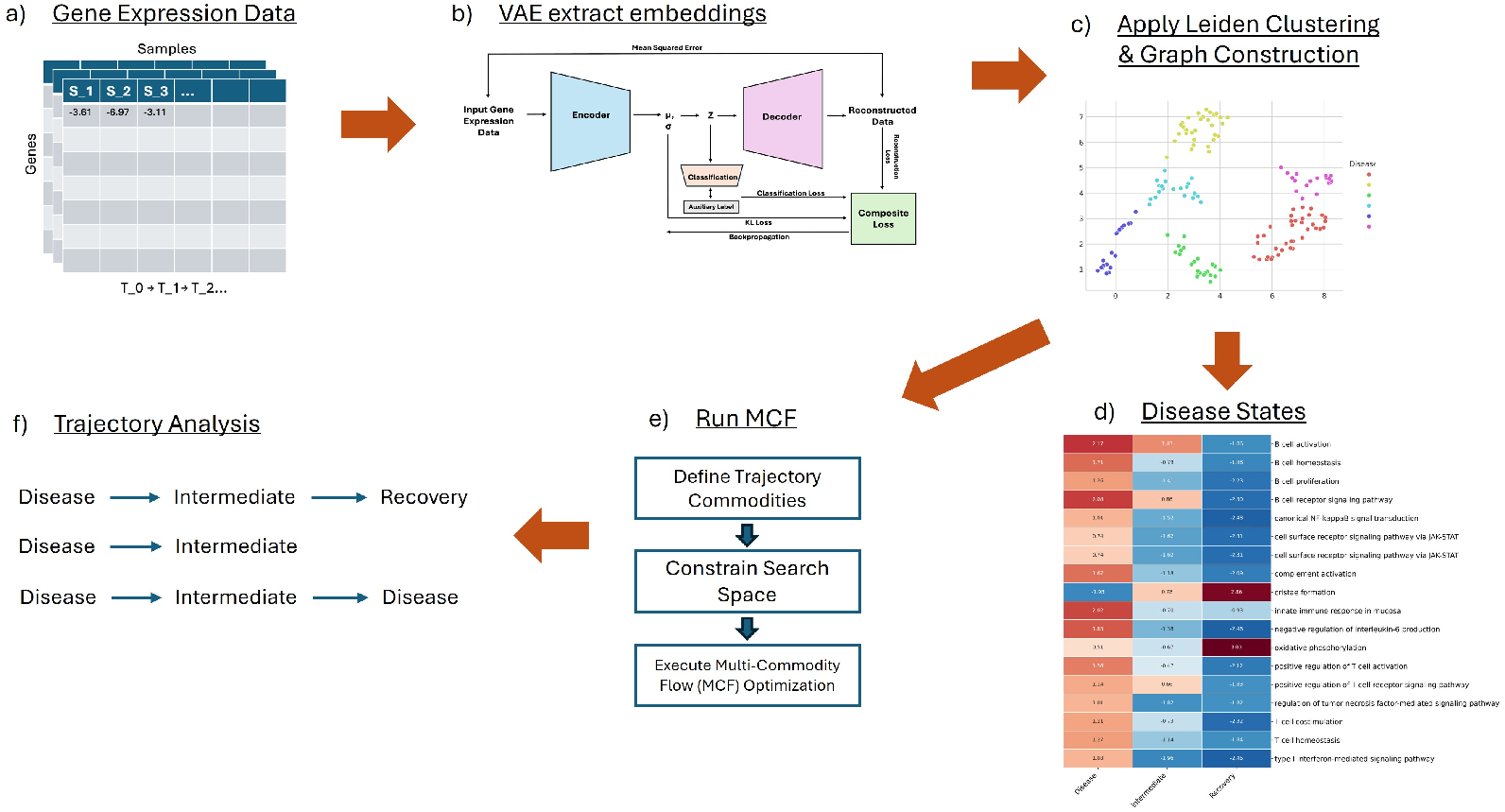
Overview of the proposed computational framework. **a)** Gene Expression Data: Input consists of longitudinal, time-series transcriptomics data from patient samples. **b)** VAE embeddings: A VAE, guided by an auxiliary classification task, learns a compressed, low-dimensional representation (embedding) of each sample. **c)** Apply Leiden Clustering & Graph Construction: The VAE embeddings are clustered using the Leiden algorithm to identify distinct patient subgroups (states), and are further constructed as a graph to run graph traversal algorithm. **d)** Biological Mechanisms: Gene Set Enrichment Analysis (GSEA) is performed on the genes of each cluster to define disease/treatment states. States were labeled as “Disease”, “Recovery”, etc., by experts. **e)** Run MCF: A MCF problem solver is formulated to model the most probable transitions between the discovered disease states over time. **f)** Trajectory Analysis: The MCF output is analyzed along with disease/treatment states to understand the key trajectories and dynamics of patient progression.

### 2.3 Sample Embedding Generation and Assessment

To learn a low-dimensional representation of patient samples, we trained a classification-guided VAE on the standardized 2000-gene expression matrix (Figure 1a,b). A classification-guided VAE was chosen over a standard unsupervised VAE to ensure the resulting latent space was not only optimized for reconstruction but was also structured by known biological labels (e.g., disease vs. healthy). This guidance (ℒ_class_) enforces a more biologically meaningful separation between key phenotypes, leading to a more robust and interpretable embedding for downstream clustering. The KL divergence ( ℒ_KL_) acts as a critical regularizer, penalizing the deviation of the learned latent distribution *q*(*z*|*x*) from a standard normal prior *p*(*z*) = 𝒩 (0, *I*). This forces the latent space to be a smooth, continuous “map” rather than a disjointed set of points, preventing overfitting and ensuring that similar patients are naturally grouped together.

The VAE architecture consisted of a fully connected encoder that compresses the 2000-gene input vector down to an intermediate hidden layer, and finally onto the target latent space. This was paired with a symmetric decoder (Figure 1b). Leaky ReLU was used as the activation function for all hidden layers. The model was trained by optimizing a composite loss function, which balanced the three objectives as a weighted sum, assuming a standard normal distribution as the prior *p*(*z*):

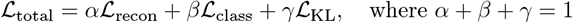

Where ℒ_recon_ is the Mean Squared Error (MSE) reconstruction loss, ℒ_class_ is the Binary Cross-Entropy (BCE) loss from a classifier head attached to the latent space, and ℒ_KL_ is the Kullback-Leibler divergence. The supervised label is either disease/healthy or treatment/pre-treatment status of patients in the dataset.

While these losses address different tasks (regression, classification, and probability matching), they can be effectively combined because ℒ_recon_ is calculated as a mean across features, bringing it to a numerical scale comparable to the other terms. The normalization constraint (*α* + *β* + *γ* = 1) ensures stable training dynamics by preventing any single loss from dominating the gradient updates. To ensure robust training and prevent overfitting, we employed k-fold cross-validation and utilized a warm-up scheme for the ℒ_class_ and ℒ_KL_ loss terms, gradually increasing their weight during the initial training epochs. The warm-up scheme stabilizes the training process, allowing the model to first learn a robust representation for reconstruction before gradually introducing the full strength of the regularization and classification guidance, which helps prevent posterior collapse [2].

To justify this multi-objective design for selecting the optimal architecture, we performed a systematic grid search evaluating both the latent space dimensionality and the different weighted composition of the loss function. We assessed performance across varying latent dimensions to determine the optimal bottleneck size for capturing biological signal without retaining noise. Concurrently, we evaluated four distinct model configurations to isolate the impact of each loss term: (1) a baseline auto-encoder (AE) optimizing only reconstruction loss ( ℒ_recon_); (2) a supervised AE adding classification guidance ( ℒ_recon_+ ℒ_class_); (3) a standard unsupervised VAE incorporating KL regularization ( ℒ_recon_+ ℒ_KL_); and (4) the proposed full model combining all three objectives (ℒ_recon_ + ℒ_class_ + ℒ_KL_). We compared these configurations using internal clustering validation metrics.

### 2.4 Disease/Treatment State Construction

To formally partition the samples, we first constructed a k-nearest-neighbor graph on the latent embeddings to build a fuzzy simplicial set, capturing the manifold’s high-dimensional structure. The Leiden community detection algorithm was then applied to this graph to partition samples into discrete clusters (Figure 1c) representing disease states. The optimal number of clusters was determined by systematically evaluating a multi-dimensional hyperparameter grid. This grid included: a) Distance Metric: metric used to construct the nearest-neighbor graph (e.g., ‘correlation’ or ‘euclidean’), b) Nearest Neighbors (*k*): The number of neighbors used to build the graph, and c) Leiden Resolution (*α*): The resolution parameter of the Leiden community detection algorithm. Each combination of these parameters yielded a different clustering partition. We selected the parameter set that produced the partition maximizing a consensus of internal validation metrics, including the Silhouette score, Calinski-Harabasz index, Davies-Bouldin index, and Correlation Difference (the average intra-cluster correlation minus the average inter-cluster correlation).

GSEA was applied on the disease/treatment states (or clusters) inferred from the clustering of sample embeddings (Figure 1d). For a given state, the gene expression values of genes in all the samples were compared to either the healthy samples (in psoriasis) or the pre-treatment samples in UC and atopic dermatitis. Gene ontology biological processes that had FDR *<* 0.05 were used to further analyze the clusters. The cluster was then characterized as “Disease” if their mechanisms indicated active disease state, “Recovery” if it indicated subduing of the pathological mechanisms, and “intermediate” if it was found to be between active pathology and recovery states. In AD, we found multiple disease states that varied in the type and granularity of immune response.

Note that a state may transcend samples of multiple patients. To understand biological mechanisms induced in the disease/treatment state, Gene Set Enrichment Analysis [18] was performed on the gene signatures of each cluster to assign meaningful biological functions. Based on expert evaluation, labels such as “disease”, “recovery”, “intermediate”, etc., were assigned to the clusters.

### 2.5 Trajectory Inference

To model patient progression through the treatment we first constructed a transition graph *G* = (*V, E*), where each node *v ∈ V* represents a sample embedding. Rather than a fully connected graph, the edges *E* were constructed by connecting each sample only to its *k*-nearest neighbors determined during the state construction step. The edge weights between a pair of samples (*u, v*) *∈ E* was calculated by taking the correlation distance (1 *− pearson*_*correlation*) of the sample embeddings. This edge weight represents the “cost” of transitioning between two biological states, penalizing transitions between dissimilar sample embeddings.

We framed the trajectory inference problem using a MCF problem solver as done in Truffle and followed the steps in Figure 1d. In this framework, each individual patient was treated as a distinct *commodity*. For a given patient, their sample at the earliest (pre-treatment) timepoint was designated as a *source* node, and their sample at the final timepoint was designated as a *sink* node. The MCF problem’s objective was to find the minimum-cost flow path for all commodities (patients) simultaneously, constrained by the temporal order (order of visits) of the samples of each patient. This process models the most likely, time-respecting progression for each patient through the network of disease states. The resulting optimal flow solution was then used to extract the dominant, time-resolved trajectory for each individual patient, mapping their unique progression through the annotated disease states.

The trajectory obtained from Figure 1d was combined with the disease/treatment states from Figure 1e to obtain potential patient trajectories in Figure 1f. Each patient’s trajectory was mapped to disease/treatment states from their first visit to the last visit, thus representing the trajectory with a set of states, *patient*_1_ = [*Disease, Intermediate, Recovery*], *patient*_2_ = [*Disease, Recovery*]. The most prominent sets of states were deemed as treatment trajectories. All treatment trajectories for the 3 datasets are shown in Figure 3.

### 2.6 Trajectory Validation and Analysis

Information from original studies (UC and psoriasis) of patients being responders or non-responders for drug treatment was used to check if the validity of the patient trajectories. The status (responder or non-responder) was determined using clinical scores in the original studies. Since patients were evaluated of being a responder or non-responder only at their last visit, the last state of a patient’s trajectory was used for evaluation. The same strategy was used to compare the method against Truffle.

## 3 Results

### 3.1 Model Evaluation

To identify the optimal architecture, we first evaluated the impact of loss function composition on cluster quality (Figure 2). A systematic grid search revealed that while classification guidance ( ℒ_class_) offered marginal improvements over a reconstruction-only baseline, integrating KL divergence ( ℒ_KL_) was critical for achieving distinct cluster separation. Consequently, the fully integrated configuration ( ℒ_recon_ + ℒ_class_ + ℒ_KL_) consistently maximized performance across all metrics, justifying its selection for downstream analysis.

**Fig. 2.**
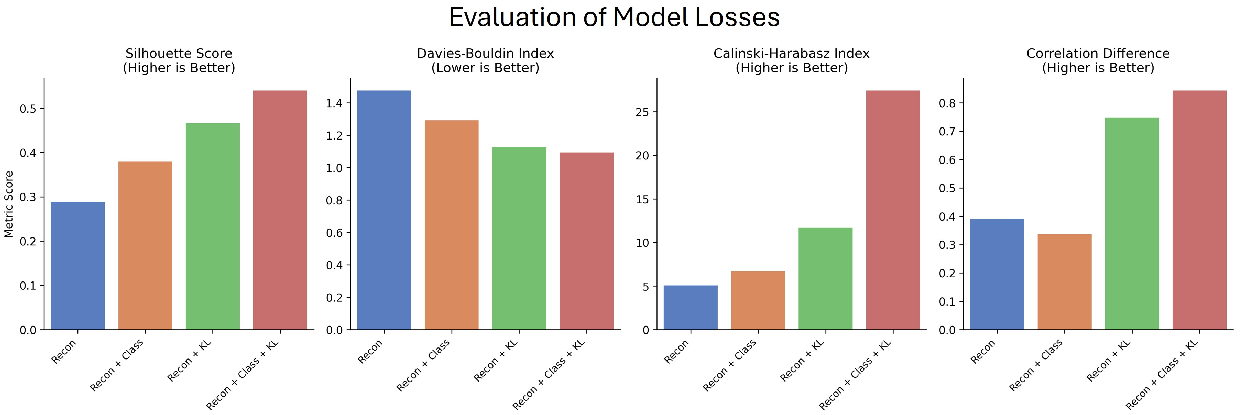
Evaluation of VAE loss function configurations.

We then benchmarked this optimized VAE against a standard PCA baseline across three datasets (UC, Psoriasis, AD). The VAE embeddings consistently outperformed PCA across four internal validation metrics. For example, in the UC dataset, the VAE achieved a Silhouette score of 0.540 (vs. 0.318 for PCA) and a Calinski-Harabasz index of 27.432 (vs. 8.331), with a significantly lower (better) Davies-Bouldin index. This confirms that the non-linear, supervised nature of our framework captures complex biological patterns more effectively than linear dimensionality reduction.

### 3.2 Dataset Findings

In the UC dataset, patients were treated with IL6 trans-signaling inhibitor. The inferred patient trajectories identified three distinct (disease/recovery/intermediate) states (Figure 3a). The first state was characterized with exclusive activation of B and T cells, the second marked a recovery state in which TNF and interleukin signaling was downregulated and recovery mechanisms such as oxidative phosphorylation and cristae formation were upregulated. In the third state B and T cell activation, JAK-STAT and NFkB signaling were downregulated and healing mechanisms such as cristae formation were upregulated. It was designated to be an “intermediate” state that indicated the patient was on the path to recovery. Overall, in the disease state immune cell activation and inflammatory pathways were upregulated, while in the intermediate and recovery states these pathways were downregulated and pathways indicating mucosal healing were upregulated. In AD, 7 states were identified that showed nuances of disease and recovery states (Figure 3b). While 3 states were detected to be disease states that varied in severity (active, disease, and mild disease), 2 were recovery (early recovery and recovery), and 2 states were close to the baseline (placebo samples - only 1 shown as unchanged). Disease states were characterized by upregulation of pathways involved in innate and adaptive immune activation as well as proinflammatory cytokine production. The different disease states potentially show the involvement of different pathogenic cell types (Th2 cells, NK cells) implicated in AD. Intermediate and recovery states were identified on the basis of downregulation of immune response and upregulation of recovery mechanisms such as T-cell homeostasis, DNA damage checkpoint signaling. While the recovery states also showed upregulation of immune activation pathways, albeit lower than disease states, they were marked by downregulation of NK and T cell activation pathways. There is a graded upregulation of TNF-mediated signaling pathway across states, with highest in the active disease state and lowest in the recovery state [6].

**Fig. 3.**
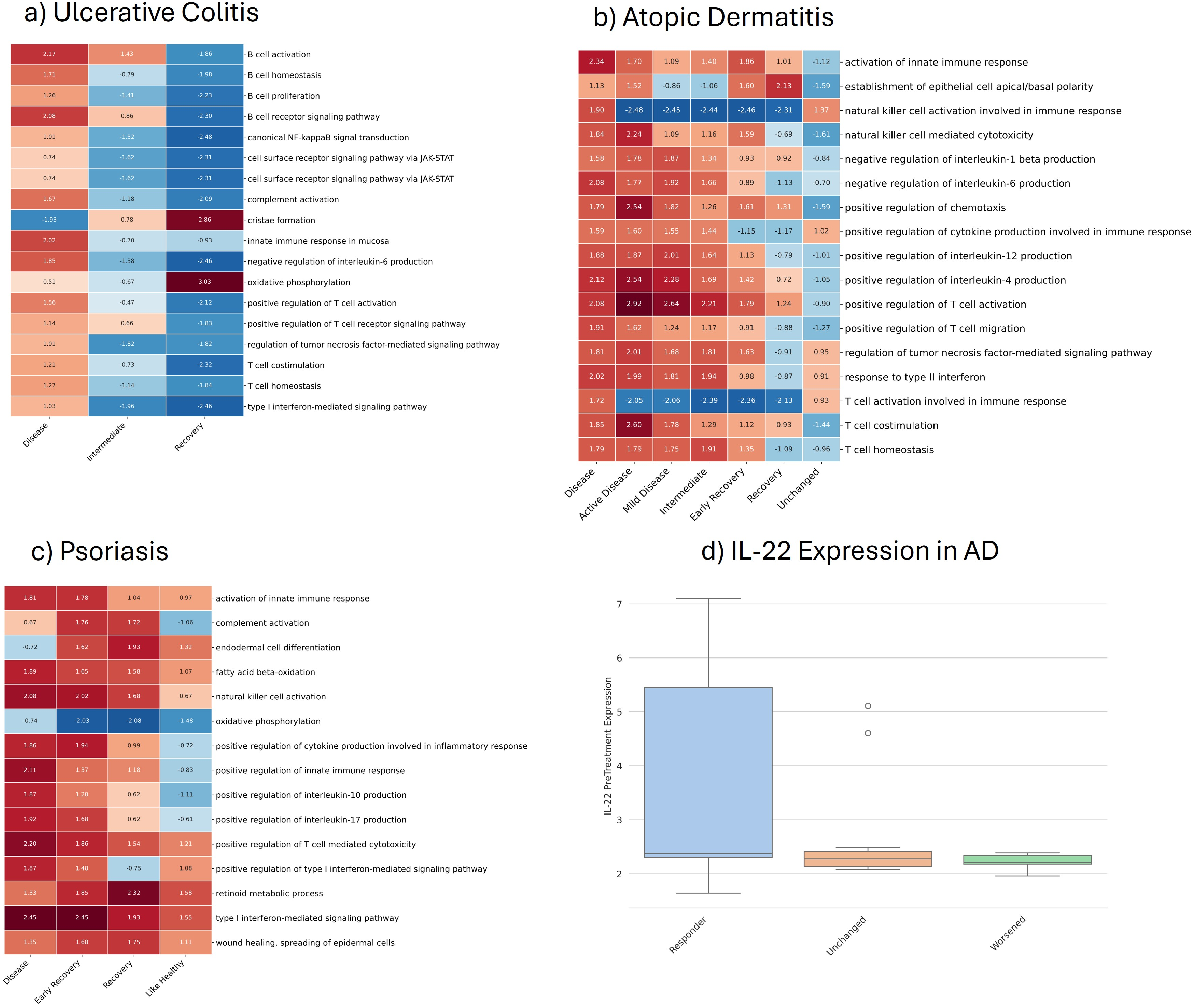
Disease/Treatment states found in the 3 datasets. NES is the normalized enrichment scores obtained from GSEA. Biological processes with FDR *<* 0.05 are shown. (a,b,c) Mechanisms in states of UC, AD, psoriasis. (d) Shows the IL-22 expression in each of the states in AD.

In the psoriasis study, we detected 4 states - disease, early recovery, recovery, and like healthy (Figure 3c). In the disease state, innate immune response, cytokine production, IL-17 pathway and cytotoxicity pathways are upregulated, whereas in the recovery state, IL-17 activity is down, and retinoid metabolic process and wound healing processes are upregulated. In the early recovery state which indicated that the patient was on their way to recovery, wound healing and retinoid pathways were also upregulated, albeit at a slightly lower level. Also, the type-1 interferon activity that is linked with psoriasis [22] has gradual down regulation from disease through like-healthy states.

### 3.3 Treatment Trajectories

Using the labels recovery/intermediate/disease of the patient states, we annotated the different states in a patient’s trajectory obtain after graph traversal (Figure 1f). Since patients were designated either as responders or non-responders depending on their clinical score at the last visit (in UC, Psoriasis), we checked if it overlaps with the last state of a patient’s trajectory. For example, if a patient was designated as a responder, we checked if their last state of the trajectory was “recovery”. Out of 9 patients in UC, our results showed 7 patients had consistent findings i.e., responders ended in “recovery” state and non-responders ended in “disease” state. In the psoriasis study, all 15 patients showed response (lower PASI scores) at the end of the 12 week treatment. Out of 15 patients, our method showed 13 patients had the recovery state as the last state in their trajectory indicating they were responders. Interestingly, 2 of the 13 responding patients showed their end state as “intermediate”.

Figure 4 lists the different trajectories that we observed across the 3 datasets. As noted earlier, in the psoriasis datasets we found 3 patient trajectories. In UC, the method detected 3 trajectories with predominant being Disease->Intermediate->Recovery and Disease->Intermediate->Disease. 2 responder patients were mapped to Disease->Intermediate trajectory. In AD, we see transitions from active-disease/disease state to mild-disease state, mild-disease to early-recovery, baseline to recovery. There are also instances of patients not changing their state through the course of treatment or getting worse as seen transitioning from mild-disease to disease and from disease to active disease. The authors in the original publication [3] noted that the remission (responders) was dependent on expression level of IL-22 as measured through microarray and qPCR, however the publication does not disclose which patients were responder and non-responders. Using the 7 disease states along trajectories of patients, we determined if a patient was a responder or non-responder (unchanged or worsened) using the following criteria: if the patient’s disease state improved then “responder”, if patient’s disease state was unchanged then “unchanged”, and if they worsened then “worsened” Figure 3d shows the expression of IL-22 gene in the 3 categories, with high expression levels for patients that were deemed responders. Patients who were unchanged or worsened had consistently low expression levels of IL-22. This confirms findings from the original publication.

**Fig. 4.**
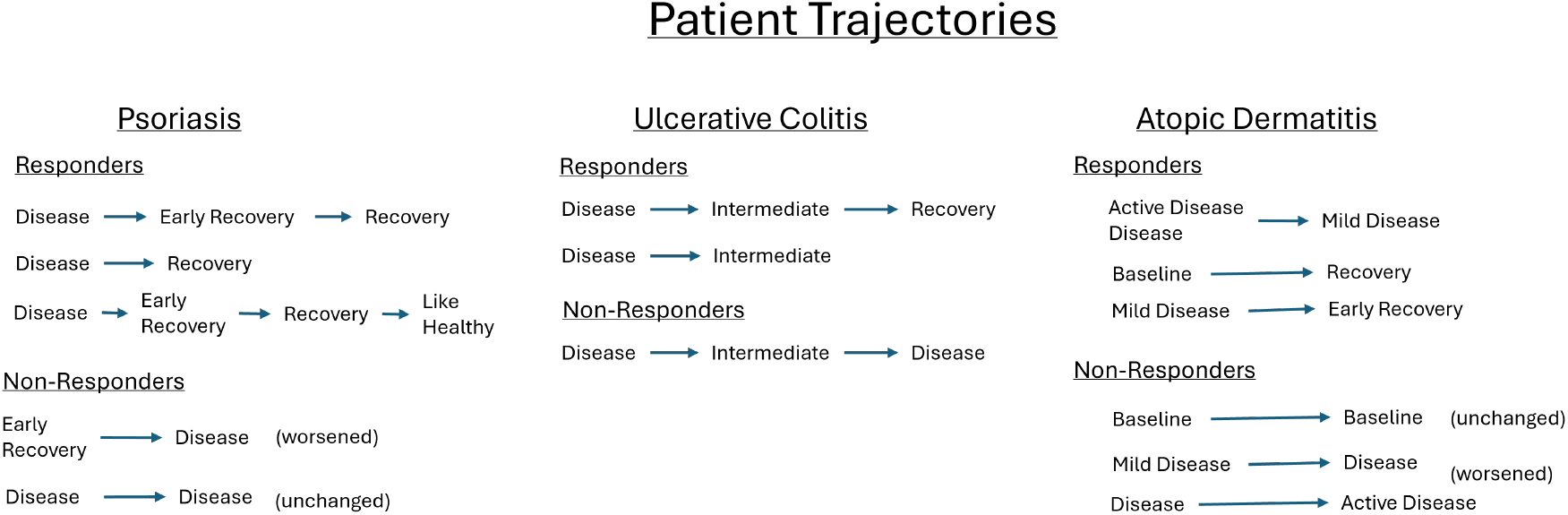
Patient Trajectories detected in the 3 datasets. States transitions are shown.

### 3.4 Comparison with Existing Approaches

Since Psupertime and Tempora only give pseudotime of samples, trajectories cannot be intuitively derived from them. Hence we compared our method to Truffle that is one of the only methods to explicitly define patient trajectories in timeseries datasets. Supplementary Figure 1 illustrates the disease states detected by Truffle in psoriasis, UC and AD (FDR *<* 0.05). In UC, mechanisms central to intervention such as IL-6 inhibition and JAK-STAT pathways [5] are not detected, and only 2 states were detected compared to 3 with our method. For the psoriasis dataset, state 2 is unrelated to psoriasis. For the AD dataset, Truffle was able to impute only 6 out of 35 patient trajectories, i.e., the trajectories for 29 patients passed through their own samples. This suggests that the Truffle was unable to leverage similar samples across patients and failed to construct dynamic disease states. For the psoriasis dataset, Truffle imputed only 4 out of 15. In contrast, our method successfully imputed 16 out of 35 AD trajectories, and 14 out of 15 trajectories in psoriasis.

Table 2 shows the number of patients detected as responders or non-responders in UC and psoriasis by the 2 methods. The # Patients column indicates the number of patients detected as responders and non-responders in the original study publications. As indicated in the methods section, the last state in the patient’s trajectory was used to check if they were responders or not and compared with the ground truth in the study datasets. Evaluation in the AD dataset was not done as patient status was not available.

**Table 2.** Comparison with Truffle.

| Dataset | # Patients | Truffle | Proposed Method |
| --- | --- | --- | --- |
| UC-Responders | 5 | 3 | 4 |
| UC-NonResponders | 4 | 1 | 3 |
| Psoriasis-Responders | 15 | 9 | 13 |

## 4 Discussion

In this study, we present a novel framework for identifying patient trajectories from longitudinal molecular data by learning latent, nonlinear representations of patient samples and constructing a trajectory for each patient. Our motivation stems from a central challenge in complex immune-mediated disorders: clinical response often manifests in gradual, heterogeneous stages rather than discrete “pre-” and “post-” treatment states. Traditional approaches such as differential expression at isolated time points or regression on primary endpoints fail to capture these intermediate disease states. As a result, potentially meaningful patterns of partial recovery or early divergence between responders can be obscured. Our method addresses this limitation by constructing latent representations of disease dynamics that emerge only when samples are modeled as trajectories rather than as independent observations.

A key component of our framework is the use of VAE to learn a continuous latent space representing disease progression. Compared with classical dimensionality reduction methods such as PCA, VAEs provide several advantages. PCA imposes a strict linear mapping that represents disease progression using global linear axes. This often collapses nonlinear structure and makes it difficult to distinguish subtle transitional states, particularly when patient recovery follows a non-monotonic pattern. In contrast, VAEs capture complex, nonlinear manifolds and preserve local neighborhood structure, allowing biologically meaningful subgroups to emerge without manual feature engineering. We evaluated the architecture using a variety of losses and size of latent representation using cluster validity metrics. Results in Figure 2 show that the fully integrated model at the optimal latent dimensions and weights of the losses achieved the highest cluster validity scores. In addition, supplementary Table 1 shows that non-linear VAE guided by the auxiliary classification task outperforms PCA in identifying tight clustering that lead to useful biological insights. These nonlinear embeddings revealed intermediate states that were not visible in the PCA space.

Biological interpretation of these trajectories provided further insight into patients with slow recovery from psoriasis. While clinical assessments frequently categorize patients as either “responders” or “non-responders,” our method identifies a continuum of partial resolution that aligns with known biology. Psoriasis treatment with an IL-17a inhibitor involves gradual down-regulation of IL-17/IL-23–associated pathways and remodeling of epidermal barrier function, both of which can occur at markedly different rates across individuals [21]. Figure 3c shows the comparative regulation of mechanisms in the 4 states detected in psoriasis, which show gradual positive regulation of healing processes, with a strong regulation in the intermediate-recovery state and then stabilizing in the recover and like-healthy states, whereas the immune response mechanisms continue to go lower in the recovery state. The disease states constructed from the latent representations captured these differences and upon analysis of patient trajectories, it revealed 2 patients with slower molecular recovery which exhibited persistent inflammatory signatures even at later time points, despite clinical improvements. Although ending molecular states of trajectories (recovery) and clinical scores were consistent for most of the patients, it is interesting to have a closer look at patients that diverge.

In UC, there were 5 responders and 4 non-responders according to the study [16]. The method correctly detected all responders, but only 2 of 4 non-responders as evaluated by the end state of the trajectory. Interestingly, the trajectory of one of the non-responding patient that was not detected by the method was predominantly in the disease state passed through the disease states before ending in an intermediate recovery state. The clinical mayo score of the patient was 3.7 which is mild-moderate disease activity (score of 0-2 is remission), pointing that the patient was on their way to recovery.

In atopic dermatitis, the method revealed 7 states of which 2 were like the baseline samples (pretreatment). Analyzing the trajectory of patients showed most of the patients with a high IL-22 expression at the pre-treatment responded to the treatment which is in line with the findings of the study [3]. Although full evaluation was not possible as the patient recovery status were not revealed, we show the IL-22 expression in different states in Figure 3d. Responder patients had higher IL-22 expression levels (mean value 3.55) compared to non-responders (unchanged 2.67, worsened 2.22). Interestingly 3 different variants of disease states were revealed (active, disease, mild).

In our search to find prior work that used longitudinal datasets to derive patient trajectories, we found only Truffle. But there was no quantitative evaluation of the inferred trajectories. Using the responder and non-responder status defined in the phase-2a studies in UC and psoriasis, we were able to objectively evaluate Truffle and our method in Table 2. Results show that our method outperforms Truffle in detection of responder and non-responder patients.

## 5 Conclusion

We developed a novel method to uncover patient trajectories in longitudinal data using non-linear latent representations of samples. The method overcomes the major bottlenecks of few timepoints and constructs a dynamic disease trajectory of patients by utilizing samples of all patients thereby revealing intermediate disease states that could not be detected in previous studies. The method was tested on 3 intervention datasets (Ulcerative Colitis, Psoriasis, Atopic Dermatitis) and results though largely agree with findings from the study publication, also show some interesting trajectory outcomes that can be further studied. One of the central assumptions of the method is that gene expression changes happen gradually over time, which is a hallmark of immune-related chronic diseases, however this assumption may not hold for diseases in the cancer domain. Also, if the patient visits are far apart, reliable reconstruction of disease dynamics may not be achieved.

## Disclosure of Interests

Funding was provided by Sanofi. Sachin Mathur, Peyman Passaban, Lu Zhang, David Habiel and Mahesh Raundhal are Sanofi employees and may hold shares and/or stock options in the company. Yixin Chen is an intern at Sanofi as of the time of submission.

All authors reviewed the article and approved it for publication.

## References

1. Angelini, F., Widera, P., Mobasheri, A., Blair, J., Struglics, A., Uebelhoer, M., Henrotin, Y., Marijnissen, A.C., Kloppenburg, M., Blanco, F.J., et al.: Osteoarthritis endotype discovery via clustering of biochemical marker data. Annals of the rheumatic diseases 81(5), 666–675 (2022)

2. Bowman, S.R., Vilnis, L., Vinyals, O., Dai, A.M., Jozefowicz, R., Bengio, S.: Generating sentences from a continuous space (2016), https://arxiv.org/abs/1511.06349

3. Brunner, P.M., Pavel, A.B., Khattri, S., Leonard, A., Malik, K., Rose, S., On, S.J., Vekaria, A.S., Traidl-Hoffmann, C., Singer, G.K., et al.: Baseline il-22 expression in patients with atopic dermatitis stratifies tissue responses to fezakinumab. Journal of Allergy and Clinical Immunology 143(1), 142–154 (2019)

4. Chen, Y., Lun, A.T., Smyth, G.K.: Differential expression analysis of complex rna-seq experiments using edger. Statistical analysis of next generation sequencing data pp. 51–74 (2014)

5. Cordes, F., Foell, D., Ding, J.N., Varga, G., Bettenworth, D.: Differential regulation of jak/stat-signaling in patients with ulcerative colitis and crohn’s disease. World journal of gastroenterology 26(28), 4055 (2020)

6. Danso, M.O., Van Drongelen, V., Mulder, A., Van Esch, J., Scott, H., Van Smeden, J., El Ghalbzouri, A., Bouwstra, J.A.: Tnf-α and th2 cytokines induce atopic dermatitis–like features on epidermal differentiation proteins and stratum corneum lipids in human skin equivalents. Journal of Investigative Dermatology 134(7), 1941–1950 (2014)

7. Hadipour, H., Liu, C., Davis, R., Cardona, S.T., Hu, P.: Deep clustering of small molecules at largescale via variational autoencoder embedding and k-means. BMC Bioinformatics 23(4), 132 (2022). https://doi.org/10.1186/s12859-022-04667-1, 10.1186/s12859-022-04667-1

8. Hasanaj, E., Mathur, S., Bar-Joseph, Z.: Integrating patients in time series clinical transcriptomics data. Bioinformatics 40(Supplement_1), i151–i159 (2024)

9. Kingma, D.P., Welling, M.: Auto-encoding variational bayes (2022), https://arxiv.org/abs/1312.6114

10. Kravitz, R.L., Duan, N., Braslow, J.: Evidence-based medicine, heterogeneity of treatment effects, and the trouble with averages. The Milbank Quarterly 82(4), 661–687 (2004)

11. Liu, J., Chang, H.W., Grewal, R., Cummins, D.D., Bui, A., Beck, K.M., Sekhon, S., Yan, D., Huang, Z.M., Schmidt, T.H., et al.: Transcriptomic profiling of plaque psoriasis and cutaneous t-cell subsets during treatment with secukinumab. JID innovations 2(3), 100094 (2022)

12. Macnair, W., Gupta, R., Claassen, M.: psupertime: supervised pseudotime analysis for time-series single-cell rna-seq data. Bioinformatics 38(Supplement_1), i290–i298 (2022)

13. Mahmud, M.S., Fu, X.: Unsupervised classification of high-dimension and low-sample data with variational autoencoder based dimensionality reduction. In: 2019 IEEE 4th International Conference on Advanced Robotics and Mechatronics (ICARM). pp. 498–503 (2019). 10.1109/ICARM.2019.8834333

14. Mathur, S., Mattoo, H., Bar-Joseph, Z.: Constrained pseudo-time ordering for clinical transcriptomics data. IEEE/ACM Transactions on Computational Biology and Bioinformatics (2024)

15. Mobasheri, A., van Spil, W.E., Budd, E., Uzieliene, I., Bernotiene, E., Bay-Jensen, A.C., Larkin, J., Levesque, M.C., Gualillo, O., Henrotin, Y.: Molecular taxonomy of osteoarthritis for patient stratification, disease management and drug development: biochemical markers associated with emerging clinical phenotypes and molecular endotypes. Current opinion in rheumatology 31(1), 80–89 (2019)

16. Schreiber, S., Aden, K., Bernardes, J.P., Conrad, C., Tran, F., Höper, H., Volk, V., Mishra, N., Blase, J.I., Nikolaus, S., et al.: Therapeutic interleukin-6 trans-signaling inhibition by olamkicept (sgp130fc) in patients with active inflammatory bowel disease. Gastroenterology 160(7), 2354–2366 (2021)

17. Smith, D.K.: Network flows: theory, algorithms, and applications. Journal of the Operational Research Society 45(11), 1340–1340 (1994)

18. Subramanian, A., Tamayo, P., Mootha, V.K., Mukherjee, S., Ebert, B.L., Gillette, M.A., Paulovich, A., Pomeroy, S.L., Golub, T.R., Lander, E.S., et al.: Gene set enrichment analysis: a knowledge-based approach for interpreting genome-wide expression profiles. Proceedings of the National Academy of Sciences 102(43), 15545–15550 (2005)

19. Tran, T.N., Bader, G.D.: Tempora: cell trajectory inference using time-series single-cell rna sequencing data. PLoS computational biology 16(9), e1008205 (2020)

20. Tyler, S.R., Chun, Y., Ribeiro, V.M., Grishina, G., Grishin, A., Hoffman, G.E., Do, A.N., Bunyavanich, S.: Merged affinity network association clustering: Joint multi-omic/clinical clustering to identify disease endotypes. Cell reports 35(2) (2021)

21. Wang, J., Ding, X.: Il-17 signaling in skin repair: safeguarding metabolic adaptation of wound epithelial cells. Signal Transduction and Targeted Therapy 7(1), 359 (2022)

22. Yao, Y., Richman, L., Morehouse, C., de los Reyes, M., Higgs, B.W., Boutrin, A., White, B., Coyle, A., Krueger, J., Kiener, P.A., et al.: Type i interferon: potential therapeutic target for psoriasis? PloS one 3(7), e2737 (2008)

23. Zhang, Y., Parmigiani, G., Johnson, W.E.: Combat-seq: batch effect adjustment for rna-seq count data. NAR genomics and bioinformatics 2(3), qaa078 (2020)

