## Supplementary for "Identification of Patient Trajectories in Timeseries Clinical Transcriptomics Data" (from another version of this preprint: version 1, 2025-12-08)

### Supplementary File - Identification of Disease Endotypes in Time-Series Clinical Transcriptomics Data

Table 1: Clustering Performance by Dataset, Distance Metric, and Method

| **Dataset** | **Distance Metric** | **Method** | Silhouette (↑) | Calinski-Harabasz (↑) | Davies-Bouldin (↓) | Corr. Diff. (↑) |
|---|---|---|---|---|---|---|
| **UC** | Correlation | VAE | 0.540 | 27.432 | 1.093 | 0.844 |
| | | PCA | 0.318 | 8.331 | 1.639 | 0.463 |
| | Euclidean | VAE | 0.283 | 26.728 | 1.099 | 0.676 |
| | | PCA | 0.163 | 8.472 | 1.901 | 0.446 |
| **Psoriasis** | Correlation | VAE | 0.511 | 31.029 | 1.253 | 0.516 |
| | | PCA | 0.427 | 15.244 | 1.455 | 0.601 |
| | Euclidean | VAE | 0.298 | 30.852 | 1.262 | 0.510 |
| | | PCA | 0.213 | 15.244 | 1.455 | 0.601 |
| **AD** | Correlation | VAE | 0.530 | 45.440 | 1.079 | 0.832 |
| | | PCA | 0.332 | 23.010 | 1.738 | 0.479 |
| | Euclidean | VAE | 0.272 | 45.493 | 1.161 | 0.780 |
| | | PCA | 0.179 | 23.010 | 1.738 | 0.479 |

*Note: (↑) Higher is better. (↓) Lower is better. Silhouette scores were calculated using the corresponding distance metric.*

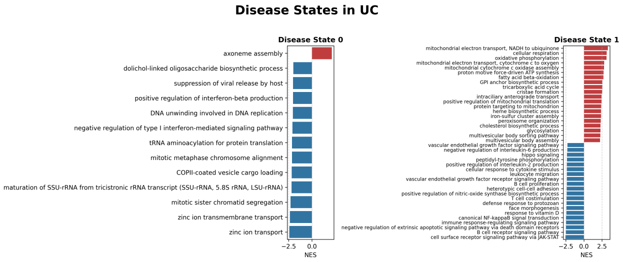


(a) GSEA on UC

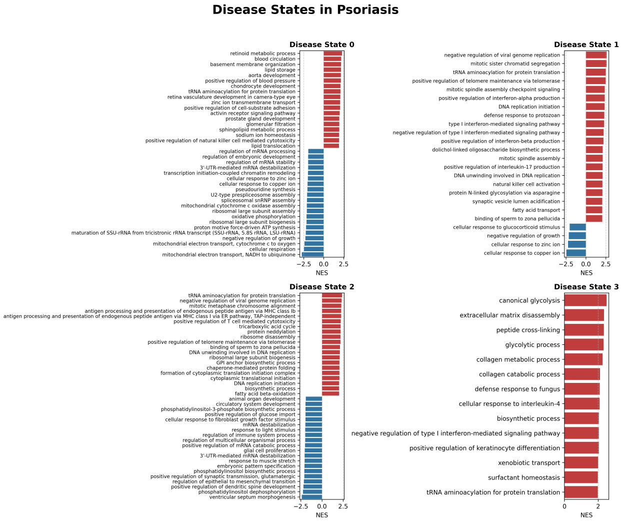


(b) GSEA on Psoriasis

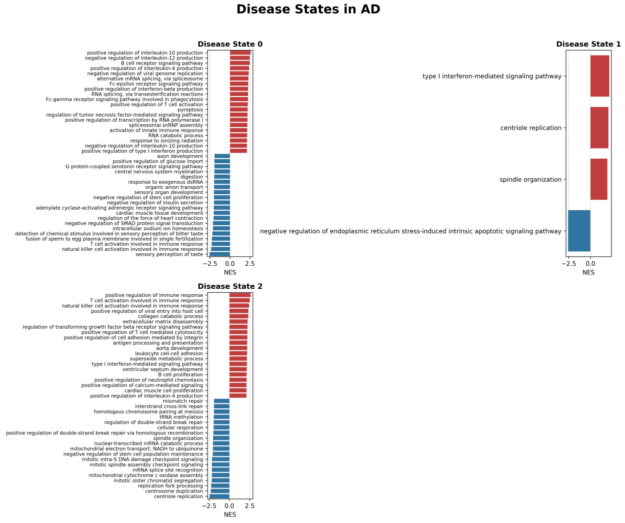


(c) GSEA on AD

Fig. 1: **GSEA for Truffle**
